## supplementary figures S1 to S11 for "Structural and molecular determinants for the interaction of ExbB from *Serratia marcescens* and HasB, a TonB paralog"

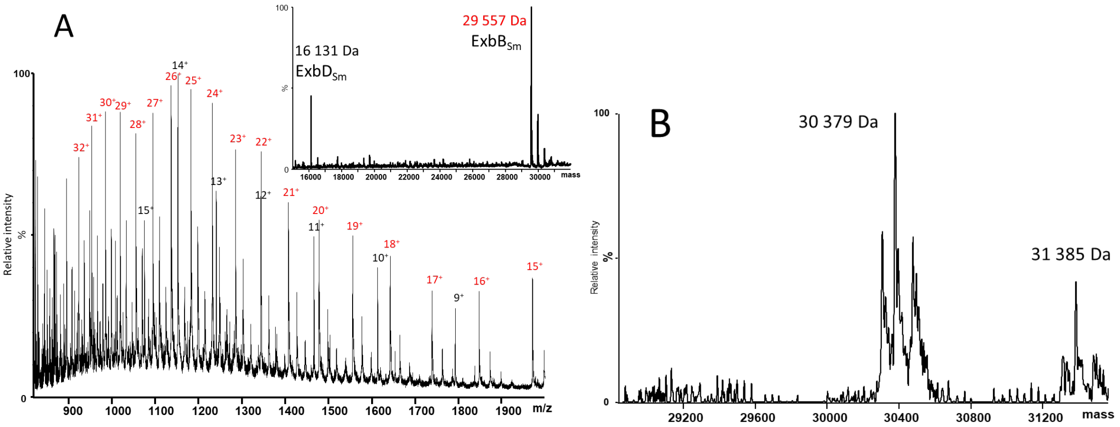


Supplementary Figure S1: Mass spectra of full-length proteins used in this study. A. MS spectrum (deconvoluted data in insert) of the ExbBD$_{Sm}$ complex in denaturing conditions (Mr$_{theoretical}$ ExbB$_{Sm}$ = 29 557 Da after signal peptide removal, Mr$_{theoretical}$ ExbD$_{Sm}$ = 16 162 Da after initial Met removal). B. Deconvoluted mass spectrum of purified ExbB$_{Sm}$ (Mr$_{theoretical}$ = 30 380 Da) showing the presence of the protein and an adduct with LMNG (Mr$_{theoretical}$ ExbB$_{Sm}$ + LMNG =31 385 Da).

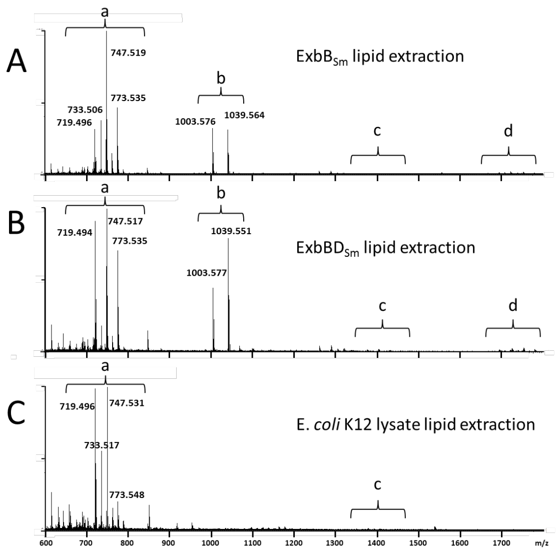


Supplementary Figure S2: Mass spectrometry analysis of lipid extracts for the following samples: A. ExbB$_{Sm}$ B. ExbBD$_{Sm}$ and C. *E. coli* lysate. The (a) area corresponds to PE (phosphatidylethanolamine) and PG (Phosphatidylglycerol) derivatives; (b) to LMNG; (c) to cardiolipins and (d) to PE/PG LMNG adducts.

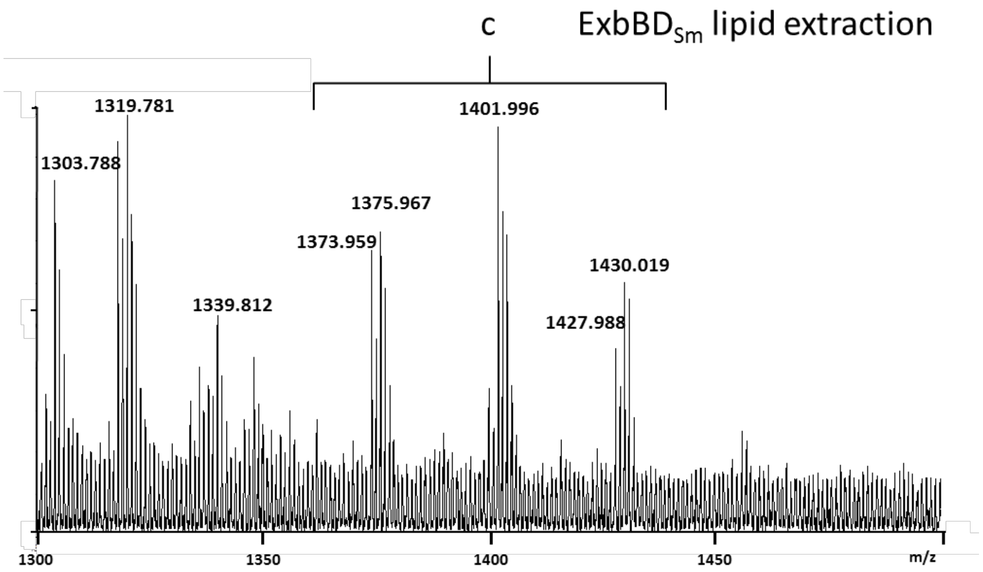


Supplementary Figure S3: Zoom of Figure S2B on the (c) area showing the different cardiolipins.

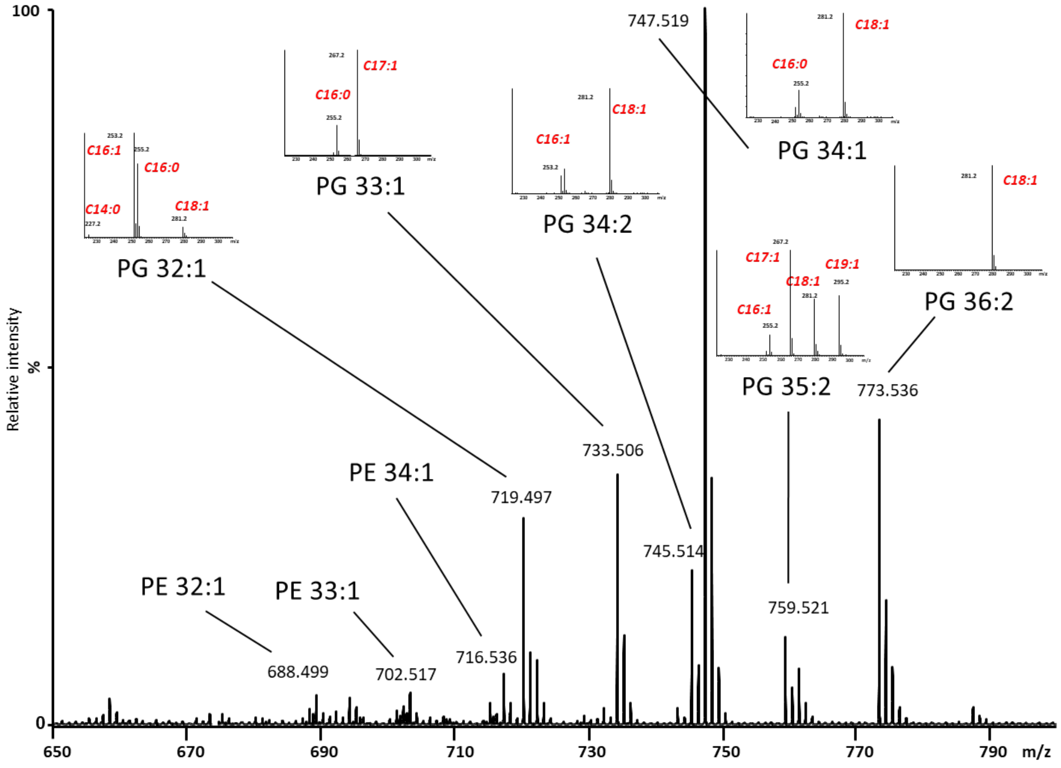


Supplementary Figure S4: Zoom of Figure S2A on the (a) area including CID spectra of the most intense ions in inserts. PE: phosphatidylethanolamine, PG: Phosphatidylglycerol, the number of carbons and insaturations are also indicated.

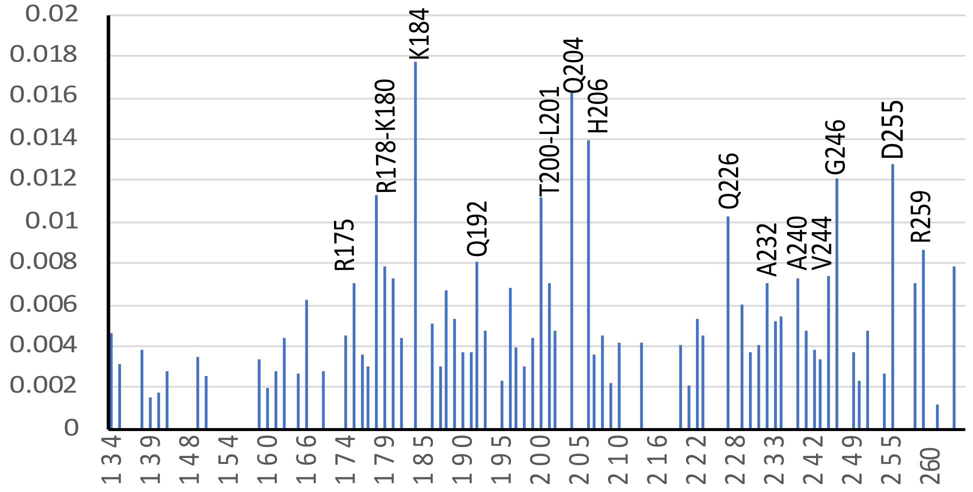


Supplementary Fig S5: Histogram showing the chemical shift perturbation (CSP) values of backbone amide signals of $HasB_{CTD}$ (0.15 mM in 50mM sodium phosphate, pH 7, 50 mM NaCl) in the presence of the $ExbB_{Sm}$ 1-44 peptide, as a function of residue numbers. The protein/peptide ratio was 1:10. Residues showing CSP higher than 0.007 are considered for analysis, except the last residue.

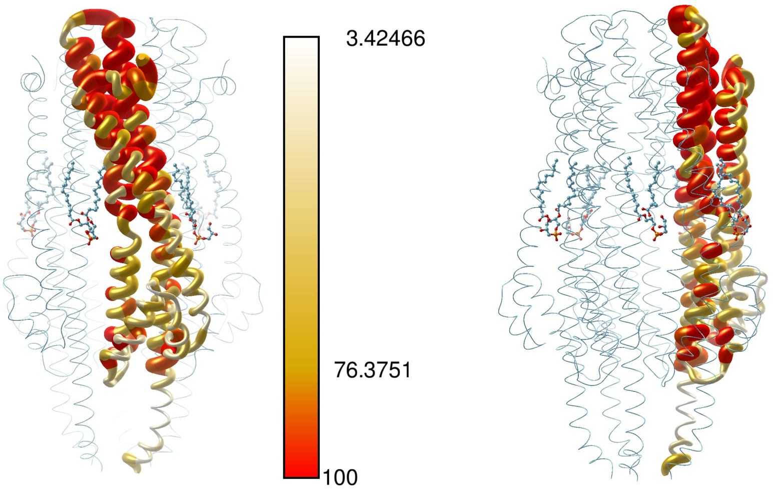


Supplementary Figure S6: sequence conservation in ExbB. structure of ExbB with one monomer ramp-colored with respect to sequence conservation using sequences retrieved in Table SI and Consurf server. Worm diameter also increases with conservation. In the transmembrane region the TM1 is the least conserved.

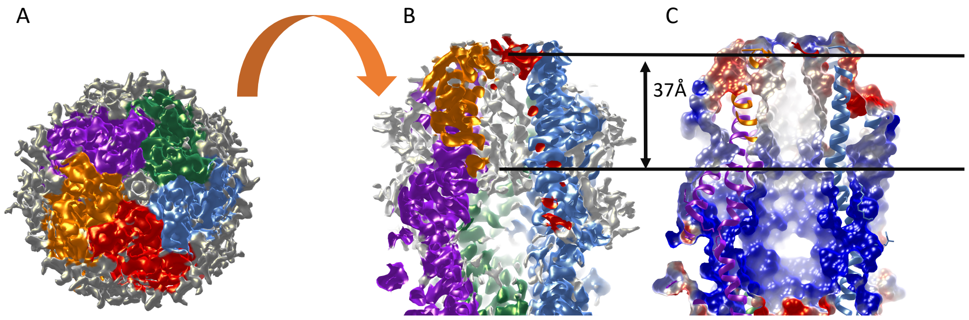


supplementary figure S7: ExbB pentamer hosts density inside its transmembrane channel. A and B, top and side views of the ExbB density colored with respect to protein chains as in Figure 8. the grey regions show extra density not accounted for by the model. C, electrostatic surface inside the channel corresponding to B. The two black lines show the cytoplasmic and periplasmic limits of the channel density. They are 37Å away from each other.

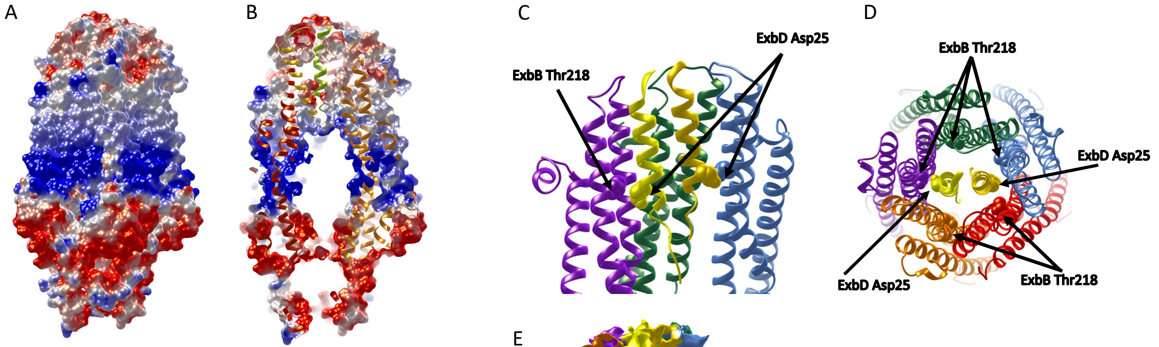


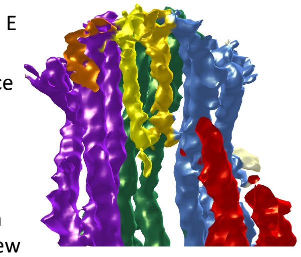


Supplementary Figure S8: electrostatic potential surface of $ExbBD_{Sm}$ determined with the APBS plugin (A, external; B, sagittal section)
Respective positions of the conserved Asp 25 residues from both ExbD TM helices and of the conserved Thr 218 residues from ExbB TM2 helices. C, sagittal section across the pentamer; D, view from the top. D, same view as C with the density map.

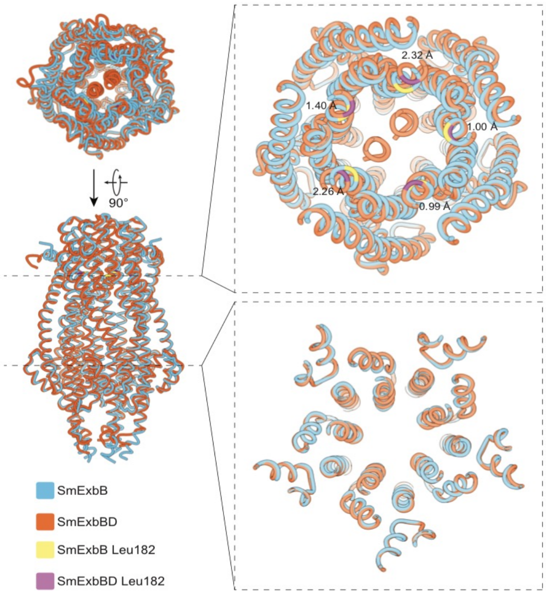


Supplementary Figure S9. Comparison of ExbBSm with ExbBDSm (top inset: section view from the periplasmic side, bottom inset: section view from the cytoplasmic side): both ExbB structures are represented in cartoon (ExbB, lightblue colour, ExbBD lightbrown colour, respectively). A specific Leu 182 residue at the periplasmic entrance is highlighted with the Ca distances between the two molecules.

A

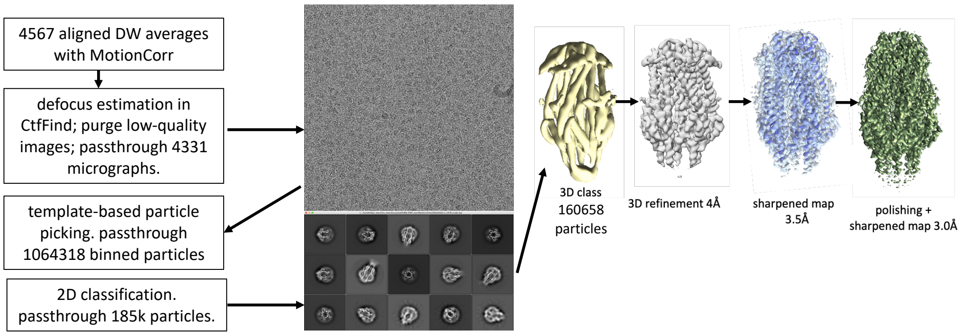


B

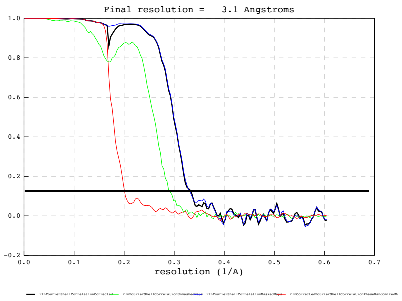


Supplementary Figure S10: Processing of ExbB cryo-EM data set (A) and final Fourier Shell Correlation plot (B).

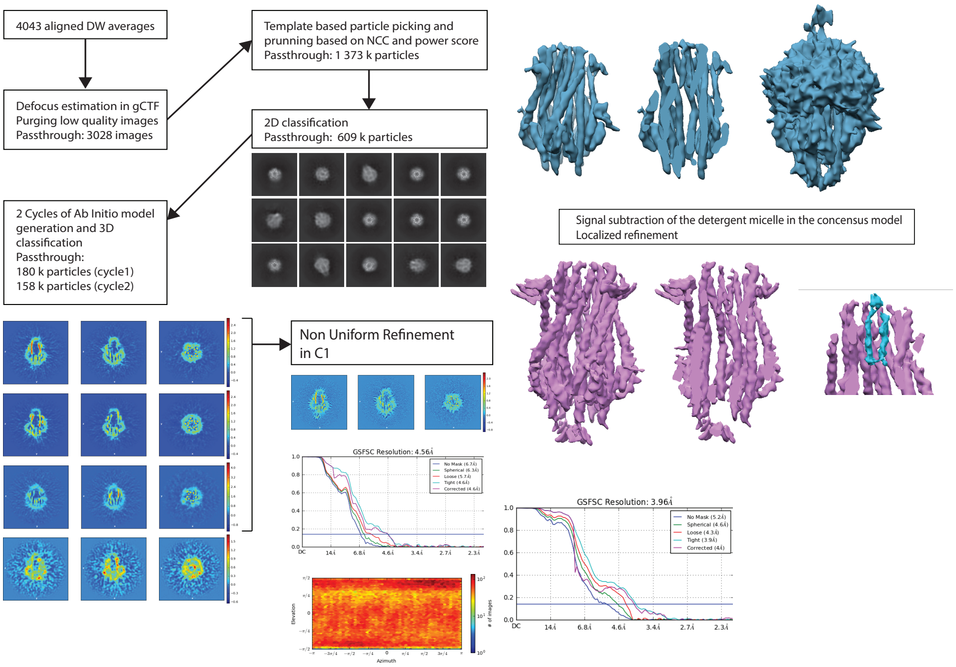


Supplementary Figure S11: Processing of ExbB-D cryo-EM data set
